# Type I PRMT inhibition protects against C9ORF72 arginine-rich dipeptide repeat toxicity

**DOI:** 10.1101/2020.05.20.106260

**Authors:** Alan S. Premasiri, Anna L. Gill, Fernando G. Vieira

**Affiliations:** ALS Therapy Development Institute, 300 Technology Square Suite 400, Cambridge, Massachusetts 02139

## Abstract

The most common genetic cause of amyotrophic lateral sclerosis (ALS) and frontotemporal dementia (FTD) is a repeat expansion mutation in the C9orf72 gene. Repeat-associated non-AUG (RAN) translation of this expansion produces five species of dipeptide repeat proteins (DRPs). The arginine containing DRPs, polyGR and polyPR, are consistently reported to be the most toxic. Here, we uncover Type I protein arginine methyltransferase (PRMT) inhibitors as possible therapeutics for polyGR- and polyPR- related toxicity. Furthermore, we reveal data that suggest that asymmetric dimethylation (ADMe) of polyGR is a determining factor in its pathogenesis.

---

A mutation in the *C9orf72* gene is the most common known cause of amyotrophic lateral sclerosis (ALS) and frontotemporal dementia (FTD)^1,2^. The mutation consists of an abnormal expansion of a repeated hexanucleotide sequence (GGGGCC) in the first intron of the *C9orf72* gene^1,2^. In ALS and FTD, the expanded nucleotide tract is translated through an unconventional mechanism known as repeat-associated non-AUG (RAN) translation^3,4^. Depending on what reading frame RAN translation takes place in, along either the sense or antisense RNA strand, it leads to the generation of five different dipeptide repeat proteins (DRPs) of variable lengths: poly-Glycine-Arginine (polyGR), poly-Proline-Arginine (polyPR), poly-Proline-Alanine (polyPA), poly-Glycine-Alanine (polyGA), and Glycine-Proline (polyGP)^3,4^.

The arginine-containing DRPs (polyGR and polyPR) in particular have been demonstrated to have detrimental effects in several model systems and to interact with several different pathways^5-7^. For example, when administered exogenously to U2OS cells, synthetic GR_20_ and PR_20_ are shown to bind to nucleoli, disrupt RNA splicing and processing, and decrease cell viability^4^. Our lab has previously demonstrated that exogenous application of synthetic GR_15_ and PR_15_ to mouse spinal cord neuroblastoma hybrid cells (NSC-34) induces cellular toxicity, as measured by various cell health and function assays, and that this toxic effect becomes more severe as the cells are further differentiated toward neurons, with primary neurons exhibiting the greatest toxicity^8^. In addition, a series of studies involving the expression of the repeat expansion in *Drosophila* have demonstrated polyGR and polyPR related toxicity^9-11^, with one study revealing the disruption of stress granule assembly due to the presence of polyGR and polyPR^11^. Other pathways that have been implicated in arginine-containing DRP toxicity include those involved in nucleocytoplasmic transport^10^ and RNA-binding^11^ though the complete nature of pathogenesis of polyGR and polyPR remains unclear. Of particular interest, recent studies in ALS suggest a role for arginine methylation in disease progression and in polyGR-related toxicity^12,13^.

PRMTs are a family of enzymes that post-translationally modify proteins by methylating nitrogen atoms of arginine residues. These modifications influence many cellular processes including transcription, RNA processing, signal transduction cascades, DNA damage response, and liquid-liquid phase separation^14^. Specifically, glycine- and arginine-rich (GAR) motifs, typical in histones and RNA binding proteins, are common targets for PRMT mediated modifications that are reported to influence protein localization and gene expression^15^. In the present study we examined whether the cytotoxic effects of exogenously applied polyGR and polyPR would be affected by pharmacological inhibition of protein arginine methyltransferase (PRMT) activity.

PRMTs are responsible for the monomethylation (MMe), asymmetric dimethylation (ADMe), and symmetric dimethylation (SDMe) of arginine residues, primarily within a GAR motif^16,17^ and are classified as Type I, Type II, or Type III depending on the type of methylation they catalyze. Type I PRMTs catalyze ADMe with MMe as an intermediate, and include PRMT1, 2, 3, 4, 6 and 8. Type II PRMTs catalyze SDMe with MMe as an intermediate, and include PRMT5 and 9. Type III PRMTs perform MMe only, and include PRMT7^18^.

To test the effects of inhibition of various PRMTs in the presence of polyGR or polyPR, we acquired commercially available, small molecule PRMT inhibitors that were capable of inhibiting either ADMe or SDMe. Because multiple PRMTs are capable of catalyzing asymmetric dimethylation of arginine residues, multiple small molecule inhibitors were selected with varying potencies against the various Type I PRMTs. These included MS023, GSK715, EPZ020411, and MS049. Because PRMT5 is the more common and abundant of the two Type II PRMTs across and in various cell types^19^, we selected a potent and specific inhibitor of PRMT5, GSK591. As a negative control, we used MS094, a previously described structural analog of MS023, which is known to be inert against all PRMT activity^20^.

We first determined the potency of the PRMT inhibitors in NSC-34 cells using an In-Cell Western (ICW) assay measuring total ADMe for Type I PRMT inhibitors, and total SDMe for the Type II PRMT inhibitor (Table 1, Fig. 1a). After establishing a working range of concentrations for each inhibitor, we measured metabolic activity (WST1 metabolism endpoint), and cytotoxicity (LDH release endpoint) in NSC-34 cells challenged by various concentrations of GR_15_ or PR_15_ with or without co-incubation with PRMT inhibitors.

**Table 1.**
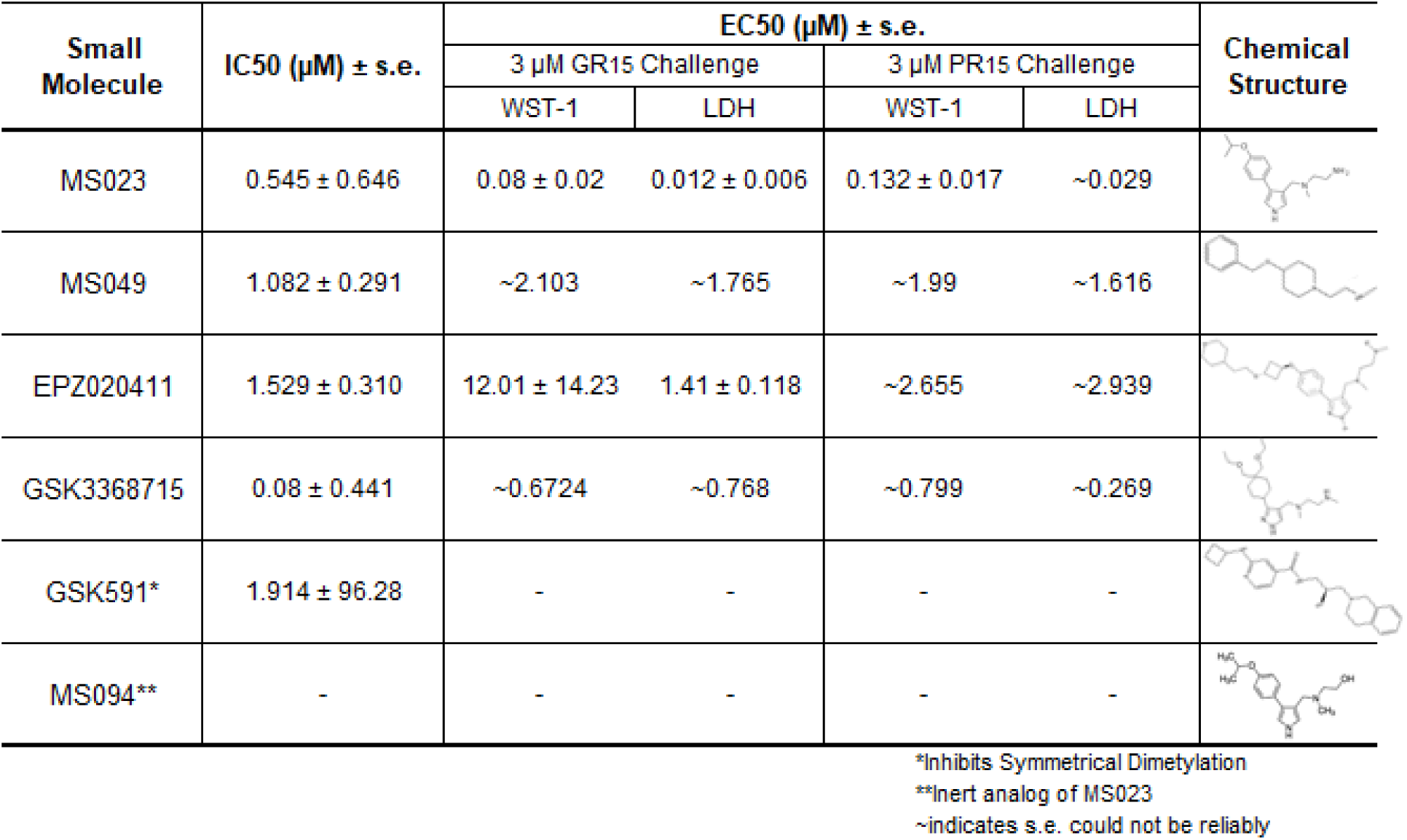
IC50s for inhibition of Dimethylation activity and EC50s for abrogation of toxicity caused by GR_15_ or PR_15_ challenge, and chemical structures for each compound tested. IC50s were calculated using a four-parameter logistical regression model and EC50s were calculated using a five-parameter logistical regression model.

**Fig. 1.**
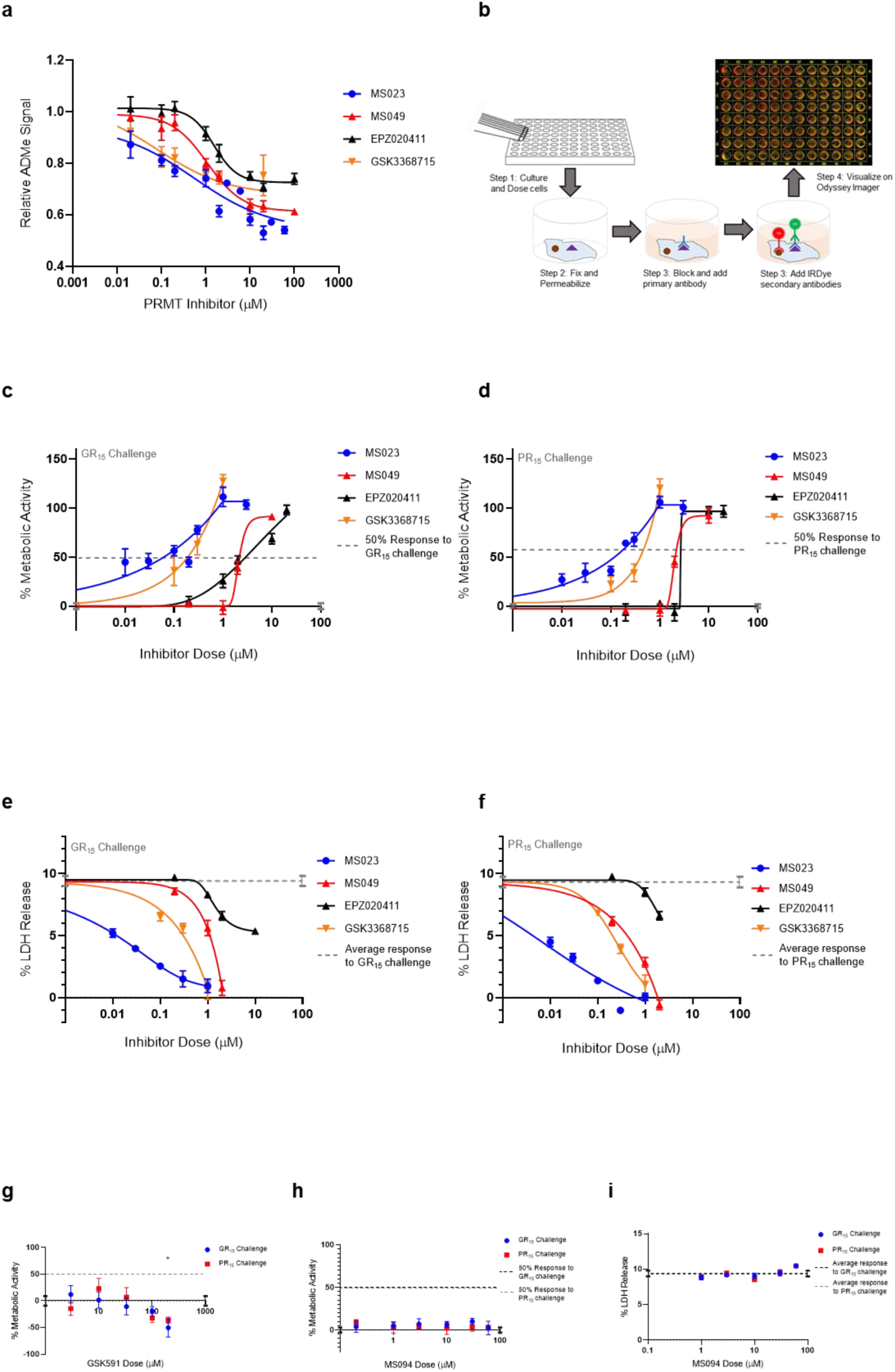
Type I PRMT inhibitors abrogate toxicity associated with GR_15_ and PR_15_ challenge in NSC-34 cells. **a**, Relative signal of total ADMe in NSC-34 cells after having been incubated with Type I PRMT inhibitors MS023, MS049, EPZ020411 and GSK3368715 for 24 hours. Quantification of the signal was done by ICW assay using an antibody against total ADMe and normalized using an antibody against total protein. IC50s were calculated using a four-parameter logistical regression model (MS023; df=132, R^2^=0.7441. MS049; df=82, R^2^=0.8218. EPZ020411; df=83, R^2^=0.7087. GSK3368715; df=50, R^2^=50). **b**, Schematic of an ICW assay workflow and example visualization. We used a primary antibody against total ADMe, and a fluorescent green IRDye secondary antibody. A red, CellTag700 antibody was used to fluorescently label total protein. **c**,**d**, Percent metabolic activity after challenging with 3 µM of GR_15_ or PR_15_ and dosing with a Type I PRMT inhibitor. We observed increases in metabolic activity by all Type I PRMT inhibitors tested with varying potencies. EC50s were calculated using a five-parameter logistical regression model (GR_15_ challenge: MS023; df=67, R^2^=0.8054. MS049; df=10, R^2^=0.9630. EPZ020411; df=13, R^2^=0.9237. GSK3368715; df=19, R^2^=0.8077; PR_15_ Challenge: MS023; df=40, R^2^=0.9034. MS049; df=10, R^2^=0.9571. EPZ020411; df=13, R^2^=0.9625. GSK3368715; df=7, R^2^=0.9330). **e**,**f**, Percent LDH release after challenging with 3 µM of GR_15_ or PR_15_ and dosing with a Type I PRMT inhibitor. We observed decreases in LDH release after dosing with the Type I PRMT inhibitors. We observed a decrease in LDH release after treatment with all Type I PRMT inhibitor tested with varying potencies. EC50s were calculated using a five-parameter logistical regression model (GR_15_ challenge: MS023; df=22, R^2^=0.9056. MS049; df=10, R^2^=0.9414. EPZ020411; df=13, R^2^=0.8807. GSK3368715; df=10, R^2^=0.9313; PR_15_ challenge: MS023; df=22, R^2^=0.9398. MS049; df=10, R^2^=0.9613. EPZ020411; df=10, R^2^=0.7078. GSK3368715; df=10, R^2^=0.9375). **g**, Percent metabolic activity after challenging NSC-34 cells with 3 µM GR_15_ or PR_15_ and dosing with GSK591. We did not observe significant abrogation of decreased metabolic activity after dosing with GSK591 (two-way ANOVA with Dunnett’s multiple comparison; n=6 untreated, n=3 treated; P values >0.1657, mean ± s.e.m). The 200 µM dose of GSK591 significantly decreased metabolic activity beyond the effect observed with 3 µM GR_15_ alone (two-way ANOVA with Sidak’s multiple comparison; n=6 untreated, n=3 treated; *P=0.0261, mean ± s.e.m.). **h**, Percent metabolic activity after challenging cells with 3 uM of GR_15_ or PR_15_ and dosing with MS094. MS094 is the inert analog of MS023, and we did not observe significant recovery of metabolic activity after dosing with MS094 (two-way ANOVA with Dunnett’s multiple comparison; NS P values >0.5496, mean ± s.e.m.). **i**, Percent LDH release after challenging cells with 3 uM of GR_15_ or PR_15_ and dosing with MS094. MS094 is the inert analog of MS023, and we did not observe decrease in LDH release after dosing with MS094 (two-way ANOVA with Dunnett’s multiple comparison; NS P values >0.1650, mean ± s.e.m.). For **c**,**d**,**g**,**h** 100% activity represents untreated NSC-34 cells, and 0% activity represents metabolic activity after 3 µM GR_15_ or PR_15_ challenge alone. For **c**,**d**,**e and f**, full dose response plots can be found in **Supplementary Fig. 1**. For **a**,**c**,**d**,**e, and f**, a full listing of *n* for each condition can be found in the **Statistics** section of the methods.

We found that the Type I PRMT inhibitors were effective at abrogating the decreased metabolic activity and increased cytotoxicity associated with the application of GR_15_ or PR_15_, with MS023 in particular demonstrating the lowest EC50s (Table 1, Fig. 1c-f). At some concentrations, incubation with Type I PRMT inhibitors resulted in complete rescue of GR_15_ or PR_15_ effects. Most of the inhibitors were inert with regards to LDH and WST1 endpoints in the absence of polyGR and polyPR at concentrations that were effective at abrogating polyGR and polyPR toxicity. The lone exception was EPZ020411. At concentrations at and above 10 µM, MS023, MS049, and EPZ020411 did reduce WST1 metabolism and elevated cytotoxicity, possibly contributing to their bell-shaped dose-response curves (Supp Fig. 1a-d, Table S1). MS094, the reported inert analog of MS023, displayed a negligible effect at inhibiting ADMe at 0.1 and 0.2 µM concentrations, and did not abrogate the GR_15_- and PR_15_-related toxicity at any concentration (Fig. 1h,i, Supp Fig. 2a). The Type II PRMT5 inhibitor, GSK591, did inhibit SDMe, but did not abrogate the decreased metabolic activity due to GR_15_ and PR_15_ challenge (Fig. 1g, Supp Fig S3). These results suggest that the activities of Type I PRMTs contribute to the toxicity produced by GR_15_ and PR_15_.

One possible interpretation of our data is that the asymmetric dimethylation of the arginine-rich DRPs is essential for arginine-rich DRP toxicity. To evaluate poly-GR as a substrate for Type I PRMT activity we conducted an in vitro methylation assay, using recombinant PRMT1 as the enzyme and S-adenosyl methionine (SAM) as the methyl donor group. We used recombinant Histone H4, a known substrate of PRMT1^26^, as a positive control for asymmetrical and symmetrical dimethylation activity. In one experiment, we used antibodies against total ADMe and Histone H4 asymmetrical dimethylation (H4R3me2a), and revealed that GR_15_ is subject to ADMe by PRMT1 in this system, and can be increasingly dimethylated when incubated with increasing amounts of PRMT1 (Fig 2a). The H4R3me2a antibody was able to detect the ADMe of GR15, possibly due to the antibody having been raised against the H4R3me2a epitope containing a methylated arginine 3, which is preceded by a glycine (Fig 2a).

**Fig. 2.**
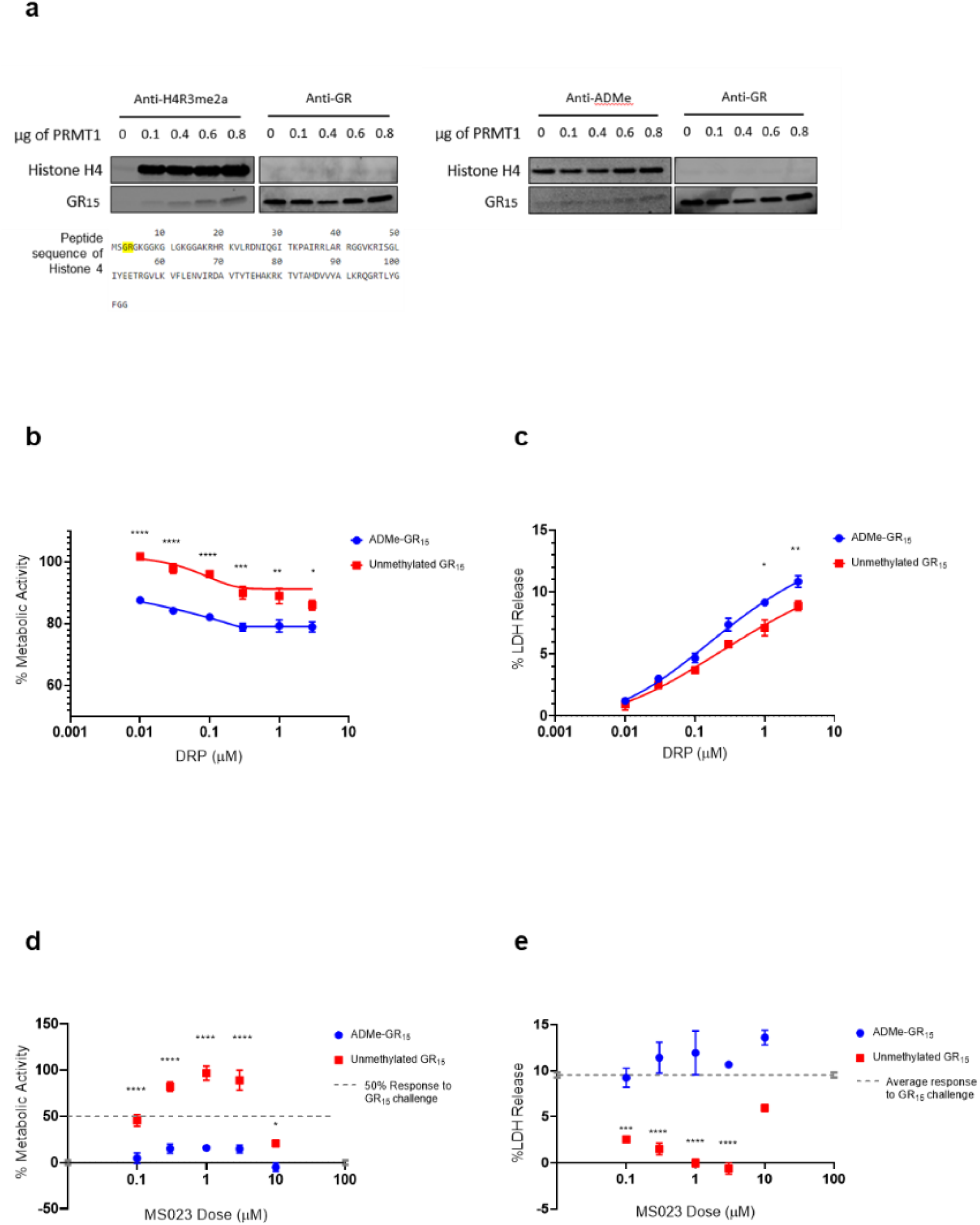
Asymmetrical Arginine Dimethylation of GR_15_ prevents abrogation of toxicity by MS023. **a**, Products of in-vitro methylation assay immunoblotted for asymmetrically dimethylated arginine 3 in Histone 4 and GR dipeptide (left) and total asymmetrical arginine dimethylation and GR dipeptide (right). Also shown is the peptide sequence of Histone 4 with the epitope of the H4R3me2a antibody highlighted. Both blots show ADMe of GR_15_ and that it is increasingly dimethylated with increasing amounts of PRMT1. **b**, Percent metabolic activity of NSC-34 cells after challenging with ADMe-GR_15_ or unmethylated GR_15_. ADMe-GR_15_ challenge caused significantly lower metabolic activity when compared to challenge by unmethylated GR_15_ (two-way ANOVA with Sidak’s multiple comparison; n=3 for each dose of DRP; ****P<0.0001, ***P=0.0002, **P=0.0013, *P=0.0281; mean±s.e.m.). **c**, Percent LDH release by NSC-34 cells after challenging with ADMe-GR_15_ or unmethylated GR_15_. Both forms of GR_15_ challenge non-significantly different levels of LDH release from 0.01 to 0.3 µM concentrations, but at higher concentrations, ADMe-GR_15_ challenge led to higher LDH release (two-way ANOVA with Sidak’s multiple comparison; n=3 for each dose of DRP; NS P>0.0593, **P=0.0083, *P=0.0117; mean±s.e.m.). **d**, Percent metabolic activity after challenging NSC-34 cells with 3 µM methylated or unmethylated GR_15_ challenge and dosing with MS023. Dosing with MS023 was not able to significantly recover the decrease in metabolic activity caused by ADMe-GR_15_ challenge, but did significantly recover metabolic activity due to unmethylated GR_15_ challenge (two-way ANOVA with Dunnett’s multiple comparison; n=9 for each dosing group; NS P>1.638, ****P<0.0001, *P=0.0411; mean±s.e.m.). **e**, Percent LDH release after challenging NSC-34 cells with 3 µM methylated or unmethylated GR_15_ challenge and dosing with MS023. Dosing with MS023 was not able to significantly reduce the LDH release caused by ADMe-GR_15_ challenge, but was able to reduce LDH release caused by unmethylated GR_15_ (two-way ANOVA with Dunnett’s multiple comparison; n=3 for each dosing group; NS P>0.0657, ****P<0.0001, ***P=0.0002; mean±s.e.m.). For **d**, 100% activity represents untreated NSC-34 cells, and 0% activity represents metabolic activity after 3 µM GR_15_ or challenge alone.

After determining that GR_15_ could be arginine methylated, we had ADMe-GR_15_ synthesized. We first compared effects of ADMe-GR_15_ challenge to effects of unmethylated GR_15_ challenge in our LDH and WST-1 assays in NSC-34 cells. ADMe-GR_15_ challenge produced similar levels of cytotoxicity as challenge with unmethylated GR_15_ peptide and caused a significant decrease in cellular metabolic function beyond the effects seen with unmethylated GR_15_ (Fig. 2b,c). To further elucidate the mechanism behind the protective effects of Type I PRMT inhibitors, we challenged NSC-34 cells with ADMe-GR_15_ and dosed with MS023. In contrast to treatment after challenge with unmethylated GR_15_, MS023 was not able to abrogate the toxicity produced by ADMe-GR_15_ challenge (Fig. 2d,e). This result suggests the toxic effects of exogenously applied GR_15_ are primarily driven by asymmetric methylation of the arginine substrates within the dipeptide repeats, rather than by aberrant methylation of endogenous proteins.

The present study reveals that Type I PRMT inhibitors can completely abrogate toxicity produced by exogenous polyGR and polyPR challenge in NSC34 cells and suggests that Type I PRMT inhibition could be a potential therapeutic strategy for C9orf72-associated ALS. We also determined that polyGR is subject to ADMe modification, and the ADMe of exogenous polyGR is crucial to the pathogenesis of the dipeptide repeat. Further studies are required to uncover if the toxic effects of polyPR are driven in a similar manner to polyGR. Additionally, experiments with different cell types, including human induced pluripotent stem cells, and DRP lengths are necessary to better understand the potential of Type I PRMT inhibitors as therapeutics for C9orf72-associated ALS and FTD. The small molecule Type I inhibitors used in the present study are each promiscuous with regards to their inhibition of various Type I PRMTs. Therefore, it is worth examining individual PRMTs to determine which ones may contribute to the effects revealed, and to what extent. In summary, our study demonstrates a novel mechanism contributing to arginine-rich DRP toxicity, and suggests a possible therapeutic strategy through Type I PRMT inhibition.

## Materials and Methods

### NSC-34 Cell Culture

NSC-34 cells (Cedarlane Laboratories, Burlington, ON, CA) were cultured in a complete medium consisting of High Glucose Dulbecco’s Modified Eagle Medium (Millipore-Sigma, Burlington, MA, USA) supplemented with 10% US-origin fetal bovine serum (Thermo-Fisher Scientific, Cambridge, MA, USA), 1% 200 mM L-Glutamine solution (Thermo-Fisher Scientific, Cambridge, MA, USA), and 1% 10,000 U/mL Penicillin-Streptomycin solution (Thermo-Fisher Scientific, Cambridge, MA, USA). Prior to preparation of NSC-34 complete medium, L-Glutamine and Penicillin-Streptomycin solutions were aliquoted and stored at −20°C, and DMEM/High Glucose was stored at 4°. At each passage, cells were washed once with Dulbecco’s Phosphate-Buffered Saline with Calcium and Magnesium (Thermo-Fisher Scientific, Cambridge, MA, USA) and treated with 0.25% Trypsin-EDTA solution (Thermo-Fisher Scientific, Cambridge, MA, USA) for 5 minutes at 37°C, 5% CO_2_ for dissociation. Prepared complete medium, DPBS, and Trypsin were always heated in a 37°C water bath before use, and stored at 4°C between uses. Cell counts for plating were performed using a Hausser Scientific cell counting chamber (Thermo-Fisher Scientific, Cambridge, MA, USA).

### Preparation of Exogenous Dipeptide Repeat Protein Solutions

Synthesized proteins GR_15_ and PR_15_ (GenicBio Limited, Kowloon, Hong Kong, CN), and ADMe-GR_15_ (Eurogentec, Leige, BE) were purchased as lyophilized powders and stored at - 20°C in a desiccator prior to reconstitution. Proteins were reconstituted in sterile DMSO (Millipore-Sigma, Burlington, MA, USA) to stock concentrations of 10 mM, and stored at 4°C.

### Preparation of PRMT Inhibitor Solutions

Small-molecule PRMT inhibitors MS023 and GSK591 (Tocris Bioscience, Briston, UK), MS049 and EPZ020411 (Cayman Chemical, Ann Arbor, MI, USA), GSK3368715 (Medchem Express, Monmouth Junction, NJ, USA), and negative control MS094 (Millipore-Sigma, Burlington, MA, USA) were purchased and stored at −20°C prior to and following reconstitution. After reconstitution using the solvents specified in Table _, stocks were aliquoted to 15-20 µL and immediately stored at −20°C. UltraPure(tm) DNase/RNase-Free Distilled Water (Thermo-Fisher Scientific, Cambridge, MA, USA

**Table _:** Small Molecule PRMT Inhibitor Details

| Drug Name(s) | Vendor Catalog # | Type of PRMT Inhibitor | Solvent Used to Reconstitute | Stock Concentration |
| --- | --- | --- | --- | --- |
| GSK591<br>GSK3203591<br>EPZ015866 | 5777/10 | Type II<br>(Symmetric) | DMSO | 10 mM |
| MS023 | 5713 | Type I<br>(Asymmetric) | Water | 10 mM |
| MS094 | SML2548 | Inactive | DMSO | 10 mM |
| MS049 | 18348 | Type I<br>(Asymmetric) | Water | 10 mM |
| EPZ020411 | 19160 | Type I<br>(Asymmetric) | DMSO | 10 mM |
| GSK715<br>GSK3368715<br>EPZ019997 | HY-128717A | Type I<br>(Asymmetric) | DMSO | 10 mM |

### In Cell Western Assay

NSC-34 cells were seeded at 3 × 10^4^ per well in clear, flat-bottom, full volume, 96 well tissue culture-treated plates (Thermo-Fisher Scientific, Cambridge, MA, USA). After plating, cells were incubated for 24h at 37°C, 5% CO_2_ prior to application of PRMT inhibitor or small molecule. Desired concentrations of compound were achieved by diluting aliquots of each stock into warmed culture medium. Cells were tested at least in quadruplicate. Once dosed, plates were incubated for 24h at 37°C, 5% CO_2_. After incubation, media was manually removed and fixed with 3.7% paraformaldehyde (Electron Microscopy Sciences, Hatfield PA, USA) in 1X PBS for 20 minutes at room temperature. Once fixed, fixing solution was removed manually and cells were permeabilized with three 0.1% Triton X-100+1X PBS (Sigma-Aldrich, St. Louis, MO, USA) washes at 5 minutes per wash. Wells were blocked using Intercept Blocking Buffer (LI-COR, Lincoln, NE, USA) for 90 minutes at room temperature with shaking. Blocking buffer was manually removed and replaced with either anti-Asymmetric Di-Methyl Arginine antibody (1:500, Cell Signaling Technology 13522) or anti-Symmetric Di-Methyl Arginine antibody (1:800 Cell Signaling Technology 13222) in Intercept Blocking Buffer and kept at 4°C overnight with no shaking. After overnight incubation, antibody solution was manually removed and washed with 0.1% Tween 20+1X PBS solution three times at five minutes per wash. An IRDye 800CW Goat anti-rabbit (1:1000, LI-COR) and CellTag700 (1:500, LI-COR) fluorescent antibody solution in Intercept Blocking Buffer was added to the plate and incubated for 60 minutes at room temperature with shaking. Antibody solution was manually removed and washed with 0.1% Tween 20+1X PBS solution three times at 5 minutes per wash. Plate was read on the LI-COR Odyssey 9120 Infrared Imaging System. Data were expressed as a ratio of 800 channel signal to 700 channel signal (test condition to total protein).

### Plating NSC-34 and Dosing with DRPs and PRMT Inhibitors

NSC-34 cells were plated at a density of 3.77 × 10^4^ cells per well in clear, flat-bottom, full volume, 96-well tissue culture-treated plates (Thermo-Fisher Scientific, Cambridge, MA, USA). One row on the top and bottom of the plate, and two columns on either side of the plate, were left without cells and contained culture medium only to minimize experimental well volume evaporation. After plating, cells were incubated for 24h at 37°C, 5% CO_2_ prior to DRP and/or PRMT inhibitor addition. At time of DRP/PRMT inhibitor addition, desired doses of DRP for challenge and inhibitor for treatment were achieved by diluting aliquots of each stock in warm culture medium. During experiments where both PRMT inhibitors and DRPs were used, PRMT inhibitors were always applied to wells first, and followed by DRP application. Vehicle controls were included as wells treated with equivalent DMSO concentrations to those that had been DRP-treated, drug treated, or both. Inhibitor toxicity controls were included as wells treated with the desired doses of drug for the experiment, but no DRP. DRP toxicity controls were included as wells only treated with the doses of DRP used for challenge. Once dosed, plates were incubated for 24h at 37°C, 5% CO_2_ prior to running the WST-1 or LDH assay endpoints. Additional controls needed for each endpoint are specified in the “WST-1 Assay” and “LDH Assay” sections of these methods.

### WST-1 Assay

Cells were plated and prepared using the steps described in the “Plating NSC-34 and Dosing with DRPs and PRMT Inhibitors” section of these methods. Other controls for this experiment included wells containing only cells in culture medium, and culture medium only. At time of testing, culture medium was removed from wells and replaced with a warmed, sterile-filtered solution consisting of DPBS with calcium and magnesium, and 4.5 g/L D-glucose (Millipore-Sigma, Burlington, MA, USA). To wells containing 200 µL DPBS-glucose solution, 20 µL/well WST-1 reagent (Millipore-Sigma, Burlington, MA) was applied and plates were then incubated at 37°C, 5% CO_2_ for 1h before plates were read at 450 nm on a SpectraMax M3 Microplate Reader (Molecular Devices, San Jose, CA, USA). As indicated in figures, WST-1 data was calculated relative to either GR/PR challenge or no challenge conditions using the following formulas. If DMSO was used as a solvent for compounds in test conditions, same concentrations of DMSO were added to Untreated Controls.

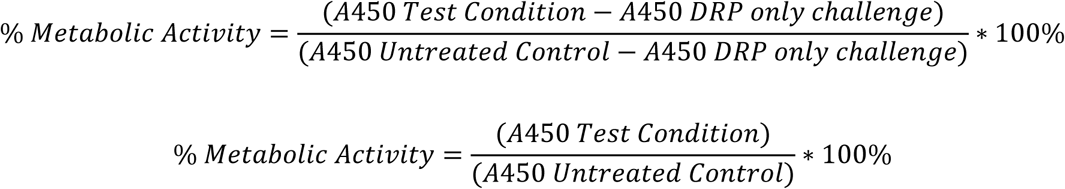

### LDH Assay

Cells were plated and prepared using the steps described in the “Plating NSC-34 and Dosing with DRPs and PRMT Inhibitors” section of these methods. Other controls for this experiment included several sets of wells with only cells in culture medium (one triplicate designated for “untreated,” one triplicate designated for “lysed” positive control), and wells with culture medium only. An additional control added only to the transfer plate at time of testing was 5 µL LDH only. Testing was performed using colorimetric LDH-Cytotoxicity Assay Kit II (Abcam, Cambridge, MA, USA) per manufacturer’s instructions. Final read at 450 nm was performed on a SpectraMax M3 Microplate Reader (Molecular Devices, San Jose, CA, USA). Data analysis included calculation of % LDH release using the following equation:

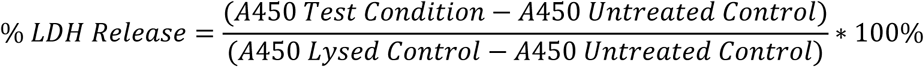

### In Vitro Methylation (IVM) Assay

Individual assays were conducted in 0.5 mL, flat cap PCR tubes (Thermo-Fisher Scientific, Cambridge, MA, USA). Systems contained Recombinant PRMT1 (Active Motif, Carlsbad, CA, USA), S-(5-Adenosyl)-L-methionine iodide (SAM, Sigma-Aldrich), either Histone H4 (Active Motif) or GR_15_ (Genic Bio), 10X PBS (Thermo-Fisher Scientific) and Nuclease-Free water (Thermo-Fisher Scientific). All reagents were added to achieve a final volume of 30 µl. Up to 0.8 µg of PRMT1, 25 µM of SAM, 3 µM of Histone H4 and 6.7 µM of GR_15_ were added to the system. 3 µl of 10X PBS was added to achieve 1X PBS. Once desired ratios of reagents were added to each tubes, systems were lightly mixed and incubated for 2h at 37°C, 5% CO_2_. After the incubation, reactions were stopped using 10 µl of 4x LDS Sample Buffer (Thermo-Fisher Scientific).

### IVM Immunoblotting

After conducting the IVM assay, samples were boiled at 95°C for 5 minutes. Samples were separated on 4-12% Bis-Tris gels (Thermo-Fisher Scientific), and blotted onto a nitrocellulose membrane. Membranes were blocked in a Superblock (Thermo-Fisher Scientific) and Tween 20 solution, then incubated with rat anti-GR (1:1000, Millipore-Sigma MABN778) and either rabbit anti-H4R3me2a (1:500, Active Motif 39006) or rabbit anti-Asymmetric Di-Methyl Arginine antibody (1:500, Cell Signaling Technology 13522). This was followed by incubation with fluorescently labelled IRDye antibodies (1:10,000 anti-rat 700 and 1:10,000 anti-rabbit 800, LI-COR 926-68076 and 925-32211) and read on the LI-COR Odyssey 9120 Infrared Imaging System. The protein standard used was the SeeBlue Plus2 Pre-stained protein Standard (Thermo-Fisher Scientific LC5925). Blots were imaged by the the LI-COR Odyssey 9120 Infrared Imaging System.

### Statistics

Statistical analyses were performed using Graphpad Prism v.8 and Microsoft Excel. Statistical tests included, two-way ANOVAs with Dunnett’s, and Sidak’s multiple comparisons tests, one-way ANOVAs with Dunnett’s multiple comparison test, five-parameter logistical regression models to calculate EC50s and a four-parameter logistical regression model to calculate IC50s. One 1 µM and 0.2 µM data point for MS049 and one 20 µM data point for EPZ020411 in **Fig 1a**. were excluded for being outside of 2 standard deviations of their respective means. A full listing of *n* values that were not represented in the figure legends are represented below.

| **FIGURE 1a.**<br>Concentration of Inhibitor (μM) | *n* | | | |
| --- | --- | --- | --- | --- |
|  | MS023 | MS049 | EPZ020411 | GSK715 |
| 100 | - | 7 | 7 | - |
| 60 | 4 | - | - | - |
| 30 | 4 | - | - | - |
| 20 | 16 | 11 | 11 | 6 |
| 10 | 20 | - | - | 6 |
| 6 | 4 | - | - | - |
| 3 | 4 | - | - | - |
| 2 | 16 | 11 | 11 | 6 |
| 1 | 20 | 11 | 11 | 6 |
| 0.2 | 16 | 11 | 11 | 6 |
| 0.1 | 16 | 4 | 4 | 6 |
| 0.02 | 16 | 4 | 4 | 6 |
| 0 | 42 | 18 | 18 | 12 |

| **FIGURE 1c,**<br>**S1a** | *n* |  |  |  |
| --- | --- | --- | --- | --- |
| Concentration<br>of Inhibitor<br>(μM) | MS023 | MS049 | EPZ020411 | GSK715 |
| 100 | - | 3 | - | - |
| 60 | 9 | - | - | - |
| 30 | 9 | - | - | - |
| 20 | - | 3 | 3 | - |
| 10 | 12 | 3 | 3 | 6 |
| 3 | 15 | - | - | 6 |
| 2 | - | 3 | 3 | - |
| 1 | 15 | 3 | 3 | 6 |
| 0.3 | 6 | - | - | 6 |
| 0.2 | 3 | 3 | 3 | - |
| 0.1 | 6 | - | - | 6 |
| 0.03 | 3 | - | - | - |
| 0.01 | 3 | - | - | - |
| 0 | 18 | 3 | 3 | 6 |

| **FIGURE 1d,**<br>**S1b** | *n* |  |  |  |
| --- | --- | --- | --- | --- |
| Concentration<br>of Inhibitor<br>(μM) | MS023 | MS049 | EPZ020411 | GSK715 |
| 100 | - | 3 | - | - |
| 60 | 6 | - | - | - |
| 30 | 6 | - | - | - |
| 20 | - | 3 | 3 | - |
| 10 | 6 | 3 | 3 | 3 |
| 3 | 9 | - | - | 3 |
| 2 | - | 3 | 3 | - |
| 1 | 9 | 3 | 3 | 3 |
| 0.3 | 3 | - | - | 3 |
| 0.2 | 3 | 3 | 3 | - |
| 0.1 | 3 | - | - | 3 |
| 0.03 | 3 | - | - | - |
| 0.01 | 3 | - | - | - |
| 0 | 12 | 3 | 3 | 3 |

| **FIGURE 1e,**<br>**S1c** | *n* |  |  |  |
| --- | --- | --- | --- | --- |
| Concentration<br>of Inhibitor<br>(μM) | MS023 | MS049 | EPZ020411 | GSK715 |
| 100 | - | 3 | - | - |
| 60 | 6 | - | - | - |
| 30 | 6 | - | - | - |
| 20 | - | 3 | 3 | - |
| 10 | 9 | 3 | 3 | 3 |
| 3 | 9 | - | - | 3 |
| 2 | - | 3 | 3 | - |
| 1 | 9 | 3 | 3 | 3 |
| 0.3 | 3 | - | - | 3 |
| 0.2 | - | 3 | 3 | - |
| 0.1 | 3 | - | - | 3 |
| 0.03 | 3 | - | - | - |
| 0.01 | 3 | - | - | - |
| 0 | 6 | 6 | 6 | 6 |

| **FIGURE 1f,**<br>**S1d** | *n* |  |  |  |
| --- | --- | --- | --- | --- |
| Concentration<br>of Inhibitor<br>(μM) | MS023 | MS049 | EPZ020411 | GSK715 |
| 100 | - | 3 | - | - |
| 60 | 6 | - | - | - |
| 30 | 6 | - | - | - |
| 20 | - | 3 | 3 | - |
| 10 | 6 | 3 | 3 | 3 |
| 3 | 6 | - | - | 3 |
| 2 | - | 3 | 3 | - |
| 1 | 9 | 3 | 3 | 3 |
| 0.3 | 6 | - | - | 3 |
| 0.2 | - | 3 | 3 | - |
| 0.1 | 3 | - | - | 3 |
| 0.03 | 3 | - | - | - |
| 0.01 | 3 | - | - | - |
| 0 | 6 | 6 | 6 | 6 |

## Acknowledgements

This work was supported by Augie’s Quest. We would like to thank people with ALS and their families and friends who have supported and inspired this work; in particular, we acknowledge those living with C9orf72 repeat expansion-mediated ALS. We would like to thank Beth Levine for her invaluable advice in the drafting of this manuscript.

## Author Contributions

Conceptualization, A.S.P., A.L.G. and F.G.V.; Data curation, A.S.P. and A.L.G.; Investigation, A.S.P and A.L.G.; Validation; A.S.P.; Methodology, A.S.P and A.L.G.; Project administration, F.G.V.; Supervision, F.G.V.; Writing—Original draft, A.S.P. and F.G.V.; Writing—Review & editing, A.L.G. and F.G.V.

