## Supplemental Material for "Type I PRMT inhibition protects against C9ORF72 arginine-rich dipeptide repeat toxicity"

### Supplementary Material

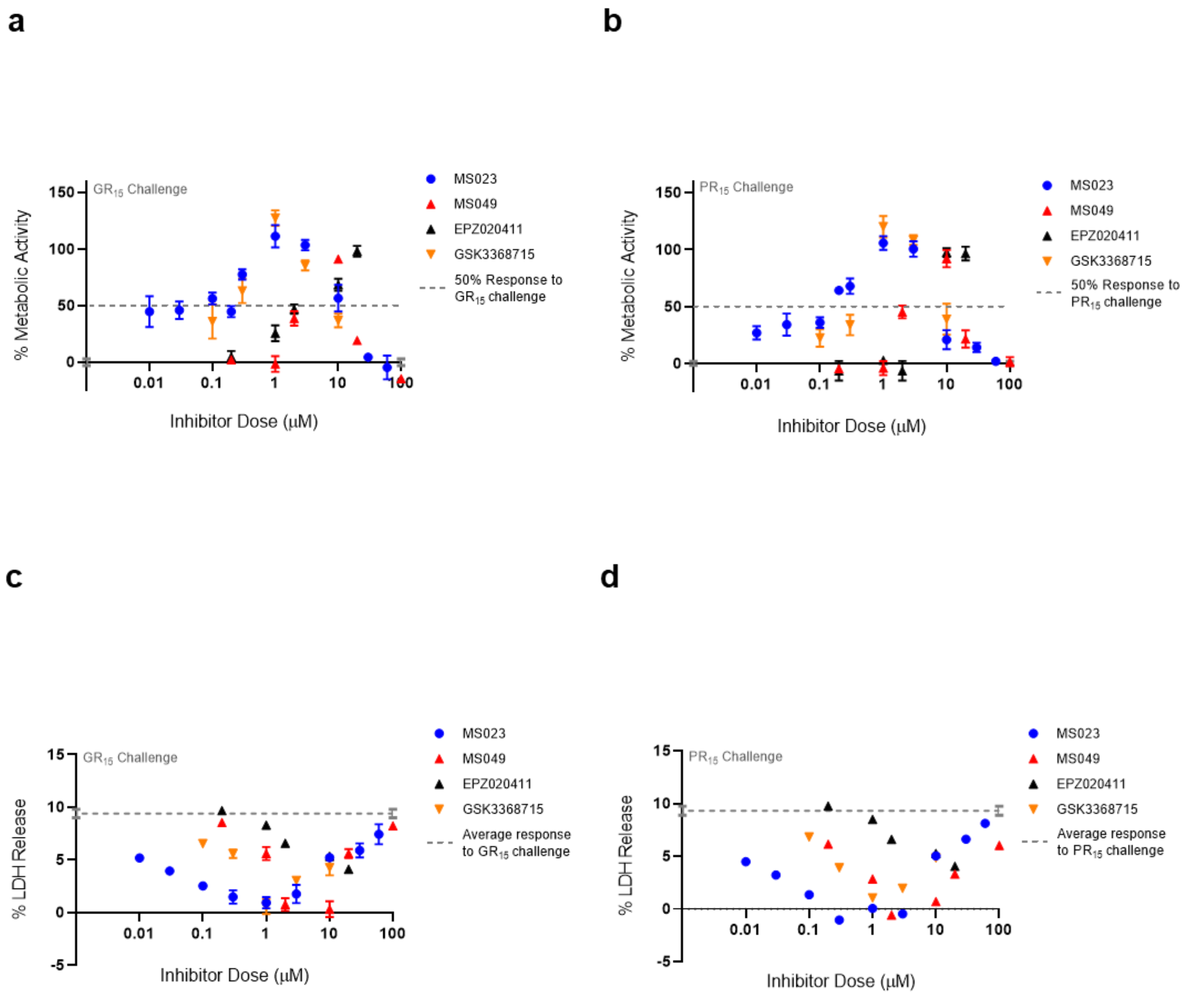


**Supplementary Figure 1**

**Figure S1: Three of four Type I PRMT inhibitor demonstrate a bell-shaped dose-response curve.**

(**a,b**) Full dose-response curves seen in **Fig 1d,e.** of percent metabolic activity after challenging NSC-34 cells with $GR^{15}$ or $PR^{15}$ and dosing with Type I PRMT inhibitors (plotted as mean±s.e.m.). (**c,d**) Full dose-response curves seen in **Fig 1f,g.** of percent LDH release after challenging NSC-34 cells with GR15 or PR15 and dosing with Type I PRMT inhibitors (plotted as mean±s.e.m.). For **a,b,** 100% activity represents untreated NSC-34 cells, and 0% activity represents metabolic activity after 3 µM $GR^{15}$ or $PR^{15}$ challenge alone. A full listing of *n* for each condition can be found in the **Statistics** section of the methods.

**a**

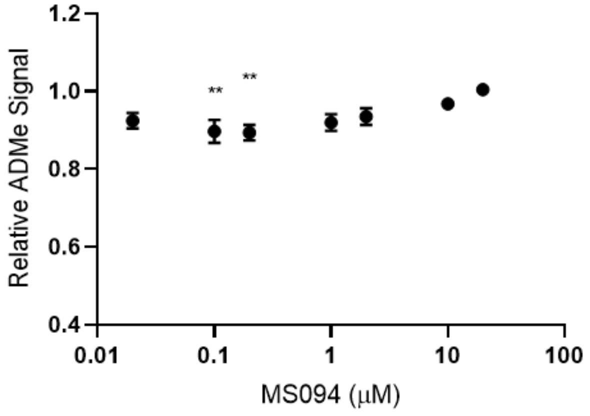


**b**

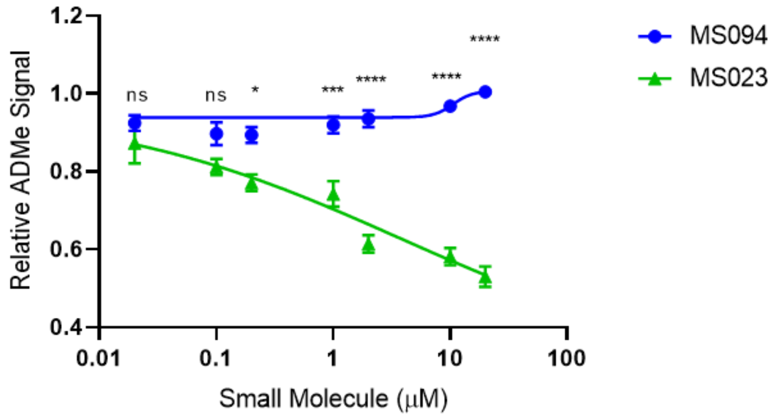


**Supplementary Figure 2**

**Figure S2: MS094 demonstrates minimal to no ability in inhibiting ADMe modification.**

(**a**) Quantified total ADMe signal in NSC-34 cells after dosing with MS094. MS094 caused significant reduction in ADMe signal at 0.1 and 0.2 μM concentrations compared to the signal of the untreated cells (one-way ANOVA with Dunnett's multiple comparison; n=12 for untreated cells, n=6 for each dosed group; NS P>0.051, control vs 0.1 μM **P=0.0059, control vs 0.2 μM **P=0.0043, mean±s.e.m.). (**b**) Compared total ADMe signal in NSC-34 cells after dosing with MS023 or MS094. MS094 demonstrates a much lower magnitude inhibition of ADMe than that caused by its active analog MS023 (two-way ANOVA with Sidak's multiple comparison; MS023 data point pulled from **Fig1a.** MS094 data points from (**a**); NS P>0.3656, ****P<0.0001, ***P=0.0003, *P=0.0471, mean±s.e.m.).

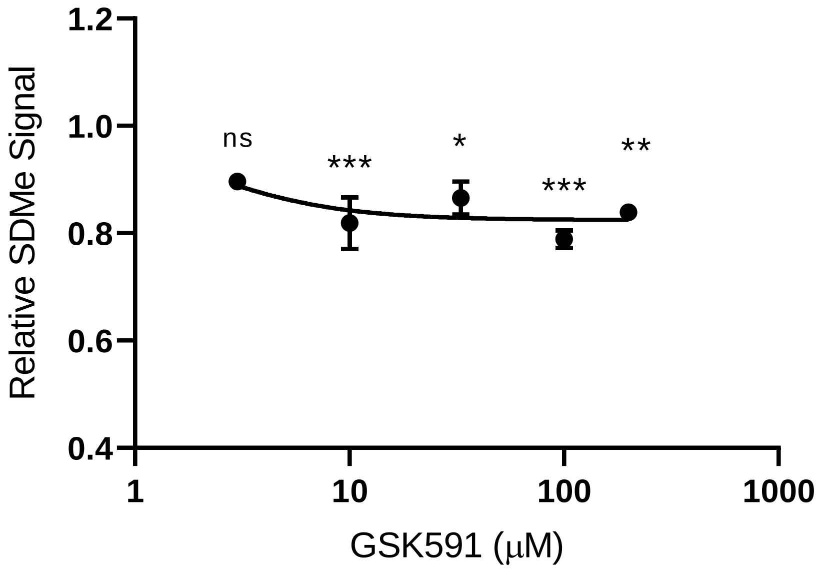


**Supplementary Figure 3**

**Figure S3: GSK591 inhibits SDMe modification in NSC-34 cells.**

Quantified total SDMe signal in NSC-34 cells after dosing with GSK591. GSK591 significantly inhibited SDMe modifications at 10 µM and above when compared to the signal of the untreated cells (one-way ANOVA with Dunnett's multiple comparison; n=10 for untreated cells, n=3 for each dosed group; NS P=0.0703, 10 µM ***P=0.0009, 100 µM ***P=0.0002, **P=0.0029, *P=0.0131, mean±s.e.m.).

**a**

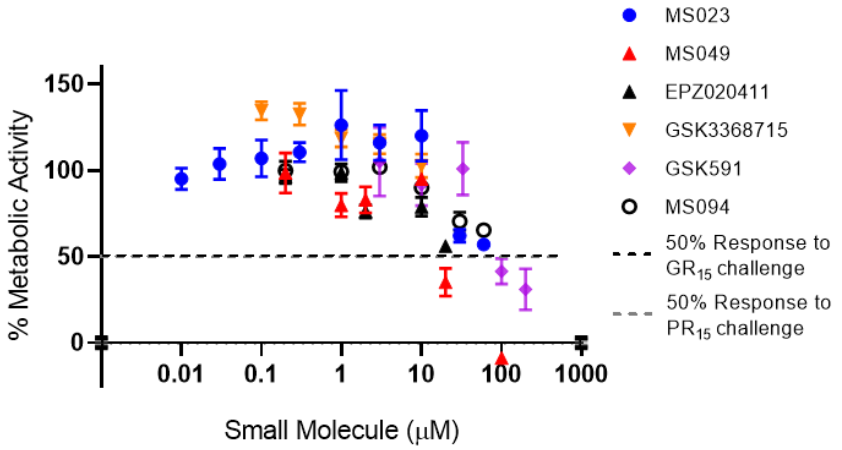


**b**

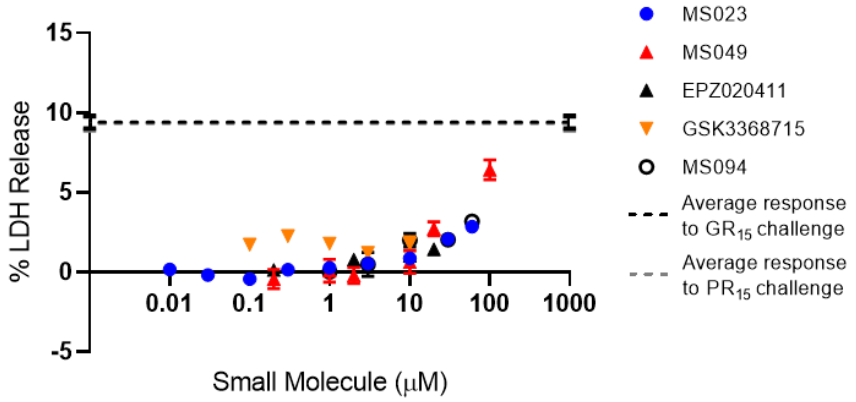


**Supplementary Figure 4**

**Figure S4: Small molecules tested exhibit some toxicity at high doses.**

(**a**) Percent metabolic activity of NSC-34 cells after applying the compounds tested. At concentrations above 10 µM, most compounds go on to show decreased metabolic activity (need table of significances?). (**b**) Percent LDH release by NSC-34 cells after applying the compounds tested. At concentrations above 10 µM, most compounds go on to an increase in

LDH release. For **a,** 100% activity represents untreated NSC-34 cells, and 0% activity represents metabolic activity after 3 µM GR15 or PR15 challenge alone. Statistical significance are represented in **Table S1**.

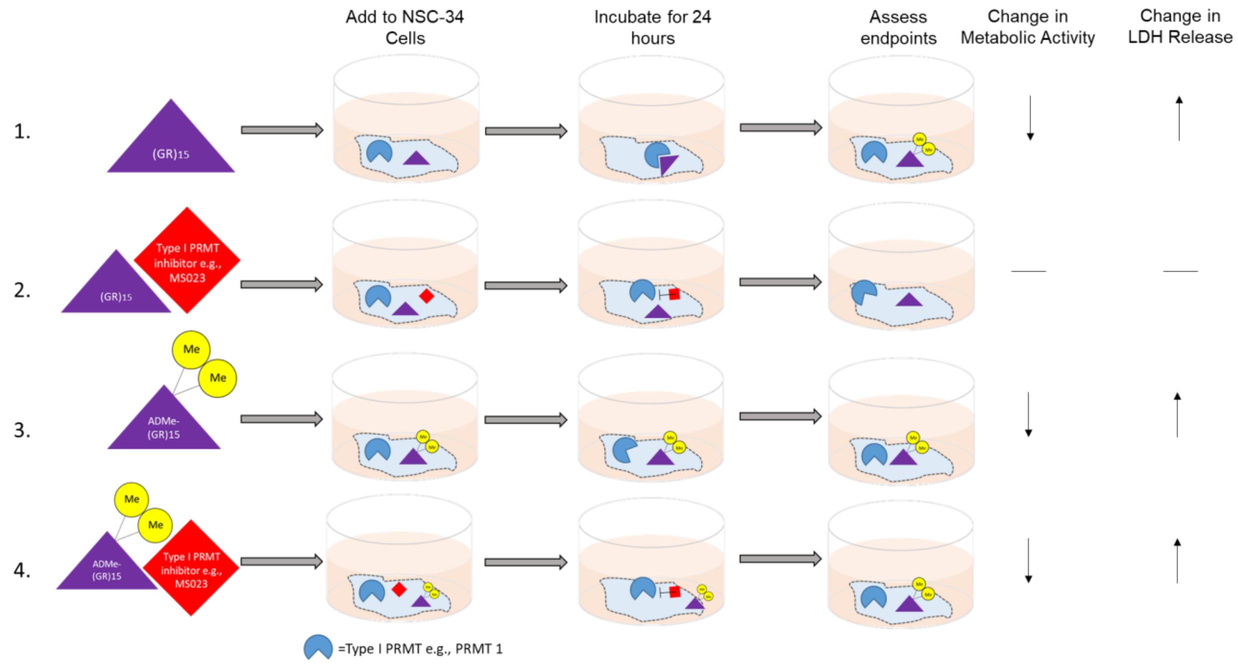


**Supplemental Figure 5**

**Figure S5: Schematic of outcomes of experiments conducted and possible mechanism of toxicity.** Based on the results in the present study the toxicity associated with $GR_{15}$ and $PR_{15}$ (not pictured) could be associated with their ability to be asymmetrically dimethylated after 24 hours of incubation (line 1). When a Type I PRMT inhibitor such as MS023 is added, the toxicity is abrogated, though the exact mechanism by which it happens remains unclear (line 2). When challenging cells with $GR_{15}$ that has been asymmetrically dimethylated, the toxic effects are still present (line 3). However when MS023 was added during the ADMe-$GR_{15}$ challenge, abrogation of toxicity was not observed, and so it is suggested that because $GR_{15}$ was already dimethylated, the PRMT inhibition had no influence on the effects seen (line 4). Taken together, the results suggest that the asymmetric dimethylation of $GR_{15}$ is the driving mechanism of toxicity seen in our assay system.

% Metabolic Activity of NSC-34 Cells after small molecule application (No GR/PR challenge)

| Dose (μM) | 200 | | 100 | | 60 | | 33 | | 30 | | 20 | | 10 | | 3 | | 2 | | 1 | | 0.3 | | 0.2 | | 0.1 | | 0.03 | | 0.01 | |
|---|---|---|---|---|---|---|---|---|---|---|---|---|---|---|---|---|---|---|---|---|---|---|---|---|---|---|---|---|---|---|
| Small Molecule | % | Sig | % | Sig | % | Sig | % | Sig | % | Sig | % | Sig | % | Sig | % | Sig | % | Sig | % | Sig | % | Sig | % | Sig | % | Sig | % | Sig | % | Sig |
| MS023 | - | - | - | - | 57.2 | * | - | - | 62.1 | ns | - | - | 120.2 | ns | 116.2 | ns | - | - | 126.3 | ns | 110.6 | ns | - | - | 107.1 | ns | 103.7 | ns | 95.2 | ns |
| MS049 | - | - | -8.8 | **** | - | - | - | - | - | - | 35.1 | **** | 95.1 | ns | - | - | 83.1 | ns | 79.9 | ns | - | - | 98.6 | ns | - | - | - | - | - | - |
| EPZ020411 | - | - | - | - | - | - | - | - | - | - | 56.2 | **** | 78.9 | * | - | - | 76.2 | ** | 98.7 | ns | - | - | 96.5 | ns | - | - | - | - | - | - |
| GSK3368715 | - | - | - | - | - | - | - | - | - | - | - | - | 102.7 | ns | 115.2 | ns | - | - | 118.6 | ns | 132.6 | ** | - | - | 134.6 | ** | - | - | - | - |
| GSK591 | 31.1 | ** | 41.3 | ** | - | - | 101.0 | ns | - | - | - | - | 91.8 | ns | 105.0 | ns | - | - | - | - | - | - | | - | - | - | - | - | - | - |
| MS094 | - | - | - | - | 65.5 | **** | - | - | 70.5 | *** | - | - | 90.1 | ns | 102.0 | ns | - | - | 99.4 | ns | - | - | 100.0 | ns | - | - | - | - | - | - |

% LDH release by NSC-34 Cells after small molecule application (No GR/PR challenge)

| Dose (μM) | 200 | | 100 | | 60 | | 33 | | 30 | | 20 | | 10 | | 3 | | 2 | | 1 | | 0.3 | | 0.2 | | 0.1 | | 0.03 | | 0.01 | |
|---|---|---|---|---|---|---|---|---|---|---|---|---|---|---|---|---|---|---|---|---|---|---|---|---|---|---|---|---|---|---|
| Small Molecule | % | Sig | % | Sig | % | Sig | % | Sig | % | Sig | % | Sig | % | Sig | % | Sig | % | Sig | % | Sig | % | Sig | % | Sig | % | Sig | % | Sig | % | Sig |
| MS023 | - | - | - | - | 2.86 | **** | - | - | 2.07 | **** | - | - | 0.87 | * | 0.54 | ns | - | - | 0.28 | ns | 0.17 | ns | - | - | -0.42 | ns | -0.17 | ns | 0.18 | ns |
| MS049 | - | - | 6.42 | **** | - | - | - | - | - | - | 2.73 | ** | 0.63 | ns | - | - | -0.21 | ns | 0.09 | ns | - | - | -0.43 | ns | - | - | - | - | - | - |
| EPZ020411 | - | - | - | - | - | - | - | - | - | - | 1.44 | **** | 1.02 | **** | - | - | 0.80 | *** | 0.46 | ns | - | - | 0.14 | ns | - | - | - | - | - | - |
| GSK3368715 | - | - | - | - | - | - | - | - | - | - | - | - | 1.77 | ** | 1.20 | * | - | - | 1.79 | ** | 2.26 | **** | - | - | 1.73 | ** | - | - | - | - |
| MS094 | - | - | - | - | 3.20 | *** | - | - | 2.04 | * | - | - | 2.01 | * | 0.50 | ns | - | - | 0.02 | * | - | - | | | - | - | - | - | - | - |

Highlighted cells indicate concentrations where the molecule showed near full abrogation of effects due to DRP challenge.

**Table S1. Effects on metabolic activity and LDH release of NSC-34 cells due to molecules in the absence of $GR_{15}$ or $PR_{15}$ challenge.** **** indicates $p<0.0001$, *** indicates $p<0.001$, ** indicates $p<0.01$, * indicates $p<0.05$. Highlighted cells represent concentrations at which molecules showed near complete abrogation of toxic effects due to $GR_{15}$ or $PR_{15}$ challenge. One-way ANOVAs with Dunnett's multiple comparisons were used to assess significance. For the WST-1 analysis, percentage of activity was compared to that after DRP challenge. 100% activity represents untreated NSC-34 cells, and 0% activity represents metabolic activity after 3 μM $GR_{15}$ or $PR_{15}$ challenge alone.
